# Dominant integration locus drives continuous diversification of plant immune receptors with exogenous domain fusions

**DOI:** 10.1101/100834

**Authors:** Paul C. Bailey, Christian Schudoma, William Jackson, Erin Baggs, Gulay Dagdas, Wilfried Haerty, Matthew Moscou, Ksenia V. Krasileva

## Abstract

**Background:** The plant immune system is innate, encoded in the germline. Using it efficiently, plants are capable of recognizing a diverse range of rapidly evolving pathogens. A recently described phenomenon shows that plant immune receptors are able to recognize pathogen effectors through the acquisition of exogenous protein domains from other plant genes.

**Results:** We showed that plant immune receptors with integrated domains are distributed unevenly across their phylogeny in grasses. Using phylogenetic analysis, we uncovered a major integration clade, whose members underwent repeated independent integration events producing diverse fusions. This clade is ancestral in grasses with members often found on syntenic chromosomes. Analyses of these fusion events revealed that homologous receptors can be fused to diverse domains. Furthermore, we discovered a 43 amino acids long motif that was associated with this dominant integration clade and was located immediately upstream of the fusion site. Sequence analysis revealed that DNA transposition and/or ectopic recombination are the most likely mechanisms of NLR-ID formation.

**Conclusions:** The identification of this subclass of plant immune receptors that is naturally adapted to new domain integration will inform biotechnological approaches for generating synthetic receptors with novel pathogen ‘baits’.

## Background

Plants have powerful defense mechanisms, which rely on an arsenal of plant immune receptors [1, 2]. Nucleotide Binding Leucine Rich Repeat (NB-LRR or NLR) proteins represent one of the major classes of plant immune receptors. Plant NLRs are modular proteins characterized by a common NB-ARC domain similar to the NACHT domain in mammalian immune receptor proteins [1]. On the population level, NLRs provide plants with sufficient diversity to maintain immunity to rapidly evolving pathogens [3, 4]. Recent findings show that novel pathogen recognition specificities can also be acquired through the fusion of non-canonical domains to NLRs [5-7] and that such fusions are widespread across flowering plants [8, 9]. These exogenous domains can serve as ‘baits’ mimicking host targets of pathogen-derived effector molecules [5, 6, 10].

The well-studied cases of NLRs with integrated domains (NLR-IDs) include *Arabidopsis thaliana RRS1-WRKY* and *Oryza sativa RGA5-HMA*. Both NLR-IDs require an additional genetically linked NLR, RPS4 and RGA4, respectively, for the activation of disease resistance [5, 10, 11]. The *RGA4/RGA5* and *RRS1/RPS4* pairs are found as neighbouring genes in inverse orientation and share a common promoter suggesting co-regulation. The products of paired NLRs form protein complexes that are essential for suppressing NLR auto-activation as well as initiation of the signaling cascade. While the NLR-ID is responsible for initial effector perception, its NLR partner is required for downstream signaling [5, 6, 10]. Whether NLR-IDs always require a genetically linked partner remains unclear.

NLR-IDs represent a successful way for plants to adopt genetic and protein linkage of NLRs to other genes to expand and diversify the pathogen recognition repertoire. On average, 10% of NLRs in sequenced plant species have been shown to contain exogenous integrated domains [8, 9]. However, little is known about the mechanisms and evolutionary history underlying NLR-ID formation.

The availability of sequenced genomes facilitates analyses of the evolution and diversification of NLR-IDs. The *Poaceae* (grasses) are a highly successful family of flowering plants that originated 120 million years ago [12, 13]. This family includes the three major cereals in modern day agriculture and human diet: maize (*Zea mays*), rice (*O. sativa*) and wheat (*Triticum* species). It has been suggested that the high genomic plasticity of grasses contributed to their adaptability and success in agriculture [14]. The genomes of sequenced grasses range in size from 270 Mb for *Brachypodium distachyon* to 17 Gb for hexaploid bread wheat (*T. aestivum*) and differ in chromosome number and ploidy level [15]. The genomes of grasses acquired diverse variation in gene copy numbers, including a high copy number of NLRs [9, 16, 17], making the *Poaceae* family an attractive system to study NLR evolution.

We examined the evolutionary dynamics of NLR-IDs in the genomes of nine grass species to address the following questions: First, were NLR-IDs distributed uniformly across subclasses of NLRs or were there specialized clades that are more prone to exogenous domain integration? Previous sequence analysis of known NLR genes, such as RGA5, hinted at the diversity of integrated domains fused to their homologs, however, no evolutionary links between these genes have been established [7, 18]. Second, we asked what was the molecular mechanism underlying NLR-ID formation.

We investigated the distribution of NLR-IDs within the NLR phylogeny and the diversity of their integrated domains within and across species. We identified several clades enriched in NLR-IDs including a monophyletic clade of NLRs that is highly amenable to repeated domain integrations from diverse gene families. The proteins within this clade showed significant lack of orthology and synteny conservation, providing evidence that orthologs acquired fusions to genes from diverse genomic locations. In addition, we identified a novel motif located upstream of integrated domains that is specifically associated with this clade and maintained across diverse NLRs. Uncovering the diversity of IDs can form the basis for new biotechnological approaches towards designing NLR receptors with synthetic fusions to new pathogen traps.

## Results

### NLR-IDs are distributed unevenly across the NLR phylogeny with one dominant clade prone to new integrations

We examined the evolution of NLRs and NLR-IDs across nine grass species with available genomes - *Setaria italica*, *Sorghum bicolor*, *Z. mays* (maize)*, B. distachyon, O. sativa* (rice)*, Hordeum vulgare* (barley)*, Aegilops tauschii, Triticum urartu*, and *Triticum aestivum* (hexaploid bread wheat). We tested two non-exclusive hypotheses about NLR-IDs:

1) The integration of exogenous domains occurs at random during NLR evolution.
2) There are conserved evolutionary integrations facilitating NLR-ID diversification.

We constructed a maximum likelihood phylogenetic tree of 4,133 NLRs from these species, based on the common NB-ARC domain. This demonstrated that while NLR-IDs occur at low frequency across the full phylogeny, a small subset of NLR clades have a much higher proportion of NLR-IDs (Figure 1A, Table 1). One major integration clade (MIC1) accounted for nearly 50% of all NLR-IDs present in the phylogeny with 59% of NLRs in MIC1 being NLR-IDs compared to 8% of proteins with NLR-IDs across all clades (Figure 1B). The MIC1 clade was found to be nested within an outer clade (Figure 1B, highlighted in blue) with only 0 to 14 % of all proteins containing NLR-IDs across the different species.

**Figure 1.**
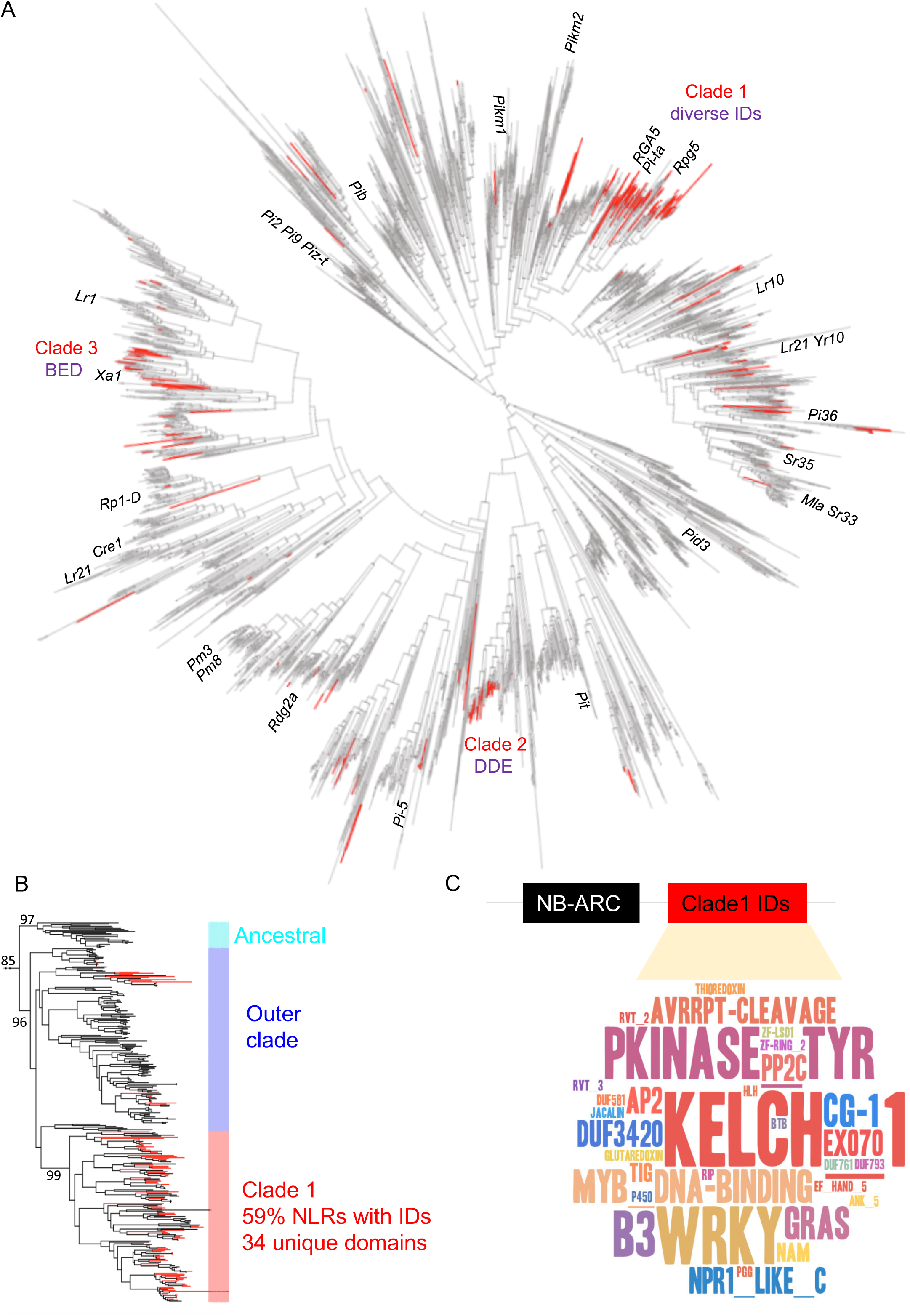
Maximum likelihood phylogeny of NLRs in grasses identifies evolutionary hotspots of NLRs with integrated domains. A) The maximum-likelihood tree of the NB-ARC family in grasses (4,133 proteins, 9 species) showing occurrence of integrated domain (ID) across the phylogeny (red branches). (B) Close-up of the MIC1 (red) as well as its outgroup clade (blue) and ancestral clade (cyan) showing the key bootstrap support values. (C) Wordcloud summary of the integrated domain diversity from MIC1. E-value cut-off for presence of an ID domain, 0.001.

**Table 1.**
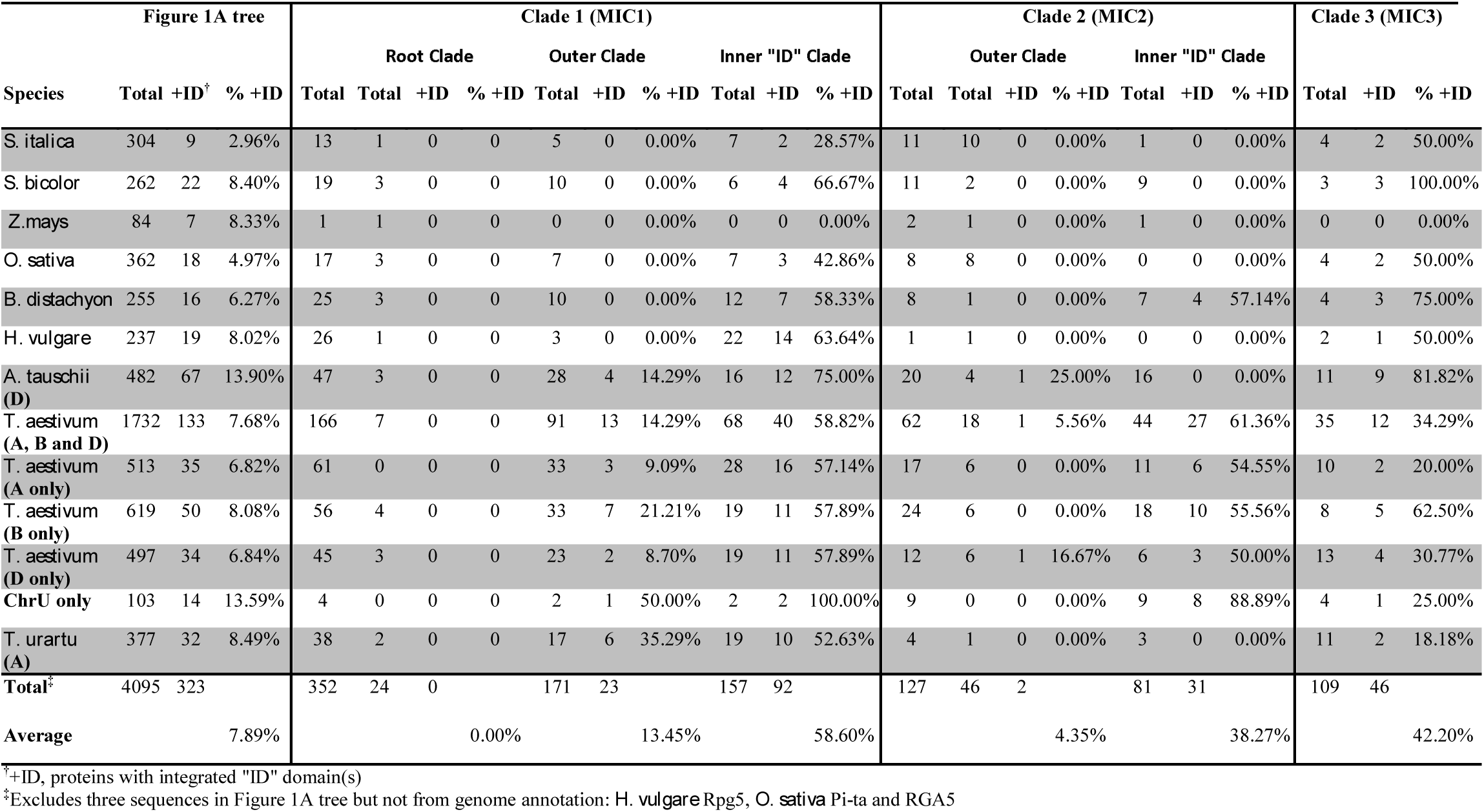
Number of NLRs and NLR-IDs in clade 1, clade 2 and clade 3 in nine grass species.

To assess whether the abundance of NLR-IDs in the major integration clades represents expansions of a single ancestral integration or repeated integrations of diverse domains we identified the domain composition within each clade. Major integration clades MIC2 and MIC3 represented expansions of ancestral integrations of DDE superfamily endonuclease and BED-type zinc finger domains, respectively (Figure 1A). In contrast, MIC1 contained a diverse range of NLR-ID proteins (Figure 1C), with integrated domains derived from 34 diverse gene families, which included both ancient and recent integrations (Figure 1C, Table 2).

**Table 2.**
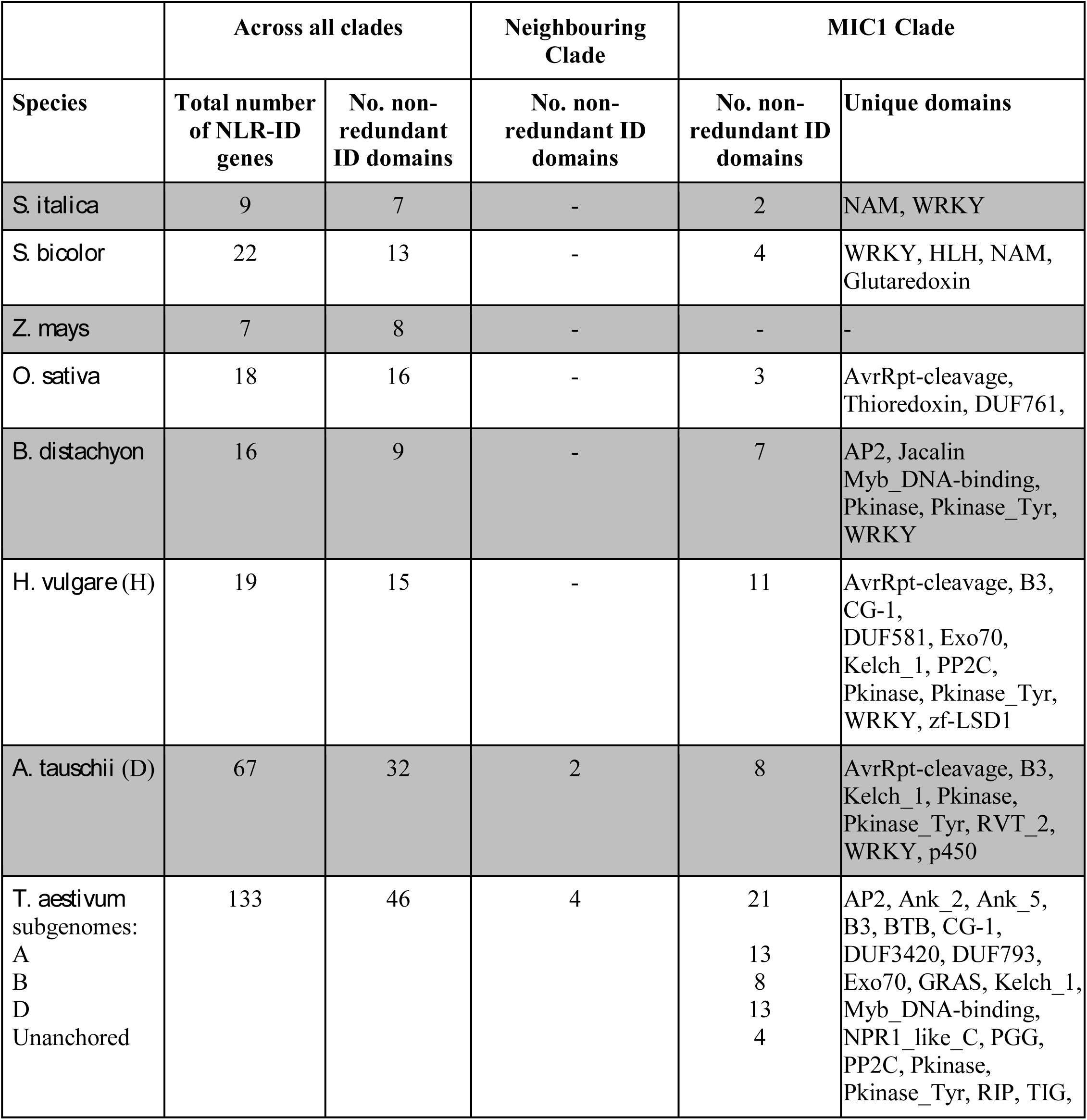
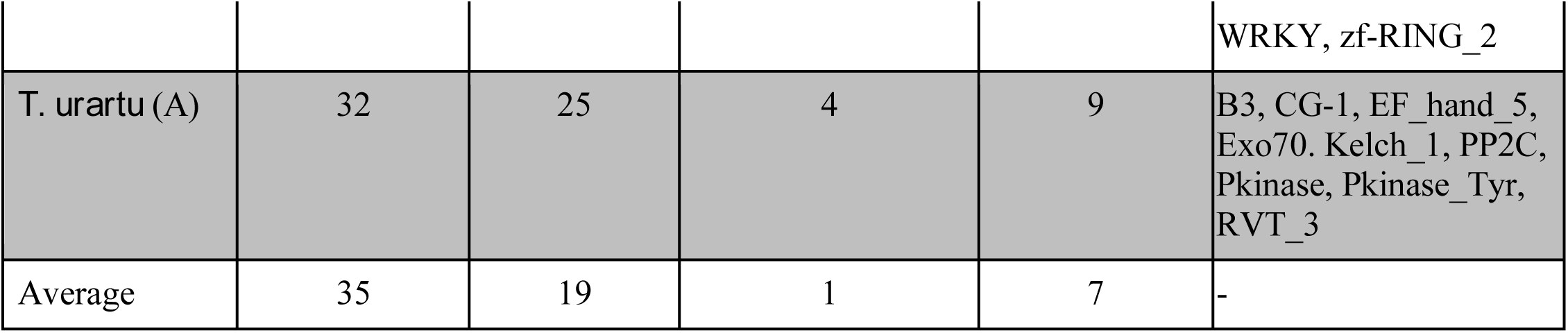
Summary of unique Pfam domains found in NLR-ID clade 1 and neighbouring clade. Only domains with e-value <1e-3 are shown. For the full list of domains with lower stringency (e-value <0.05), see Additional file 14

|  | Across all clades |  | Neighbouring Clade | MIC1 Clade |  |
| --- | --- | --- | --- | --- | --- |
| Species | Total number of NLR-ID genes | No. non-redundant ID domains | No. non-redundant ID domains | No. non-redundant ID domains | Unique domains |
| *S. italica* | 9 | 7 | - | 2 | NAM, WRKY |
| *S. bicolor* | 22 | 13 | - | 4 | WRKY, HLH, NAM, Glutaredoxin |
| *Z. mays* | 7 | 8 | - | - | - |
| *O. sativa* | 18 | 16 | - | 3 | AvrRpt-cleavage, Thioredoxin, DUF761, |
| *B. distachyon* | 16 | 9 | - | 7 | AP2, Jacalin Myb_DNA-binding, Pkinase, Pkinase_Tyr, WRKY |
| *H. vulgare* (H) | 19 | 15 | - | 11 | AvrRpt-cleavage, B3, CG-1, DUF581, Exo70, Kelch_1, PP2C, Pkinase, Pkinase_Tyr, WRKY, zf-LSD1 |
| *A. tauschii* (D) | 67 | 32 | 2 | 8 | AvrRpt-cleavage, B3, Kelch_1, Pkinase, Pkinase_Tyr, RVT_2, WRKY, p450 |
| *T. aestivum* subgenomes:<br>A<br>B<br>D<br>Unanchored | 133 | 46 | 4 | 21<br>13<br>8<br>13<br>4 | AP2, Ank_2, Ank_5, B3, BTB, CG-1, DUF3420, DUF793, Exo70, GRAS, Kelch_1, Myb_DNA-binding, NPR1_like_C, PGG, PP2C, Pkinase, Pkinase_Tyr, RIP, TIG, |
|  |  |  |  |  | WRKY, zf-RING_2 |
| *T. urartu* (A) | 32 | 25 | 4 | 9 | B3, CG-1, EF_hand_5, Exo70, Kelch_1, PP2C, Pkinase, Pkinase_Tyr, RVT_3 |
| Average | 35 | 19 | 1 | 7 | - |

We included 38 well-studied NLRs from monocots into the phylogeny to see which of those are contained in MIC1 (Figure 1A). Known resistance genes within MIC1 included *RGA5*, *Rpg5* and *Pi-ta*, which encode NLR-HMA, NLR-kinase and NLR-thioredoxin, respectively [10, 19-22].

### Proliferation of MIC1 in grasses is accompanied with continued domain shuffling

We examined the composition of NLR-IDs in MIC1 for each of the nine grass genomes under study (Figure 2A, Additional file 1). As diverse NLR-IDs were present in all the studied grass species with the exception of *Z. mays*, we predict that this clade originated before the split of Panicodae, Ehrhartoidae and Pooidae at least 60 MYA [23]. Following the evolution of the *Poaceae* genomes, MIC1 seems to have expanded after the Brachypodium/Triticeae split with a substantial increase in NLR-IDs occurring in the allohexaploid bread wheat genome. The MIC1 clade includes 68 proteins (40 with IDs) and IDs contributed by 21 different gene families. However, the relative ratio of NLRs with and without extra domains in this clade has remained relatively constant at around 59% suggesting a stable rate of domain integration across these species (Table 1).

**Figure 2.**
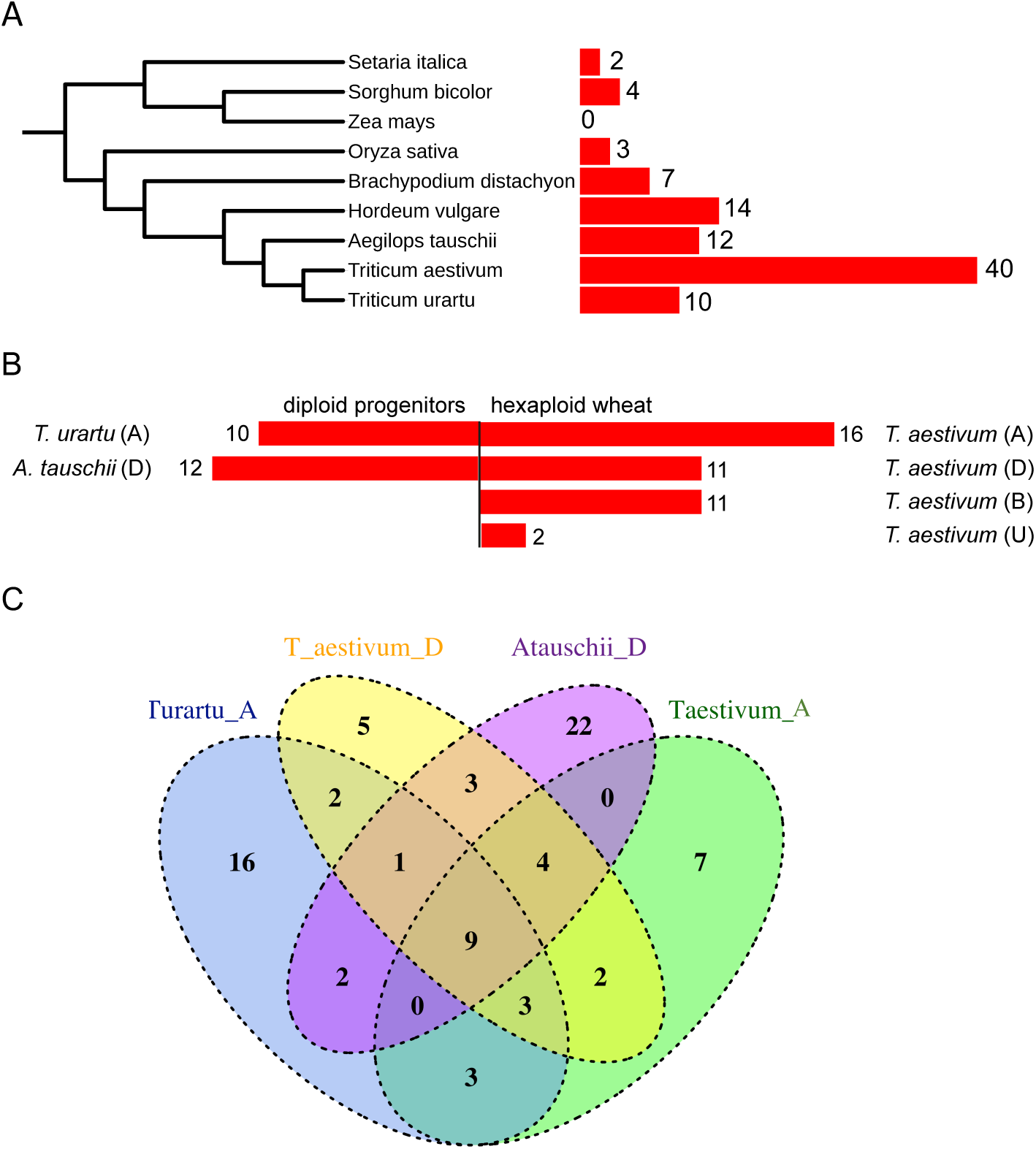
MIC1 has proliferated in grasses and continues to accumulate new domains as seen from comparison of wheat and its diploid progenitors. (A) Overall evolutionary relationship of grasses used in this study and corresponding number of NLR-IDs in MIC1. (B) Wheat sub-genomes contain a similar number of NLR-IDs in MIC1 as their diploid progenitors. (C) The repertoires of IDs are different among wheat sub-genomes and their progenitors suggesting continuous integration of new IDs.

The allohexaploid nature of the wheat genome (A, B, D genomes) and availability of genomes from two diploid progenitors (A, *T. urartu* and D, *A. tauschii*) allowed us to test if the increase in NLR-IDs in wheat represents a simple addition of diploid genomes or if new integrations are continuing to occur. We found that the numbers of NLR-IDs in the A and D genomes of *T. aestivum* to *T. urartu* and *A. tauschii* were highly similar (Figure 2B). However, the domain compositions across NLR-IDs from the A, B, D genomes of wheat and the A and D diploid progenitors did not overlap completely, indicating a continuous integration of new domains after the divergence of these species (Figure 2C). It is possible that differences in the observed repertoires can be explained partly by incomplete genome annotations or fragmented assembly of NLRs. However, the A, B and D subgenomes of wheat are of the same quality and contain full-length NLRs [24]. This suggests that the observed differences cannot be just explained by varying assembly quality. In our later analyses (see results section, “Duplication of genes encoding IDs followed by translocation of either ID or NLR lead to new NLR-ID formation”) we were able to identify homoeologous genes within wheat that contained distinct IDs. Moreover, continued integrations are supported by the overall trend in expansion of ID repertoires across the grass genomes. The genome assemblies of *B. distachyon*, *Z. mays* and *O. sativa* are of much higher quality than those of the *Triticeae* species, yet they contain fewer NLR-IDs and have lower ID diversity.

### NLR-IDs in MIC1 form genetic pairs with NLRs from another clade

Since two of well-studied NLR-IDs require a genetically linked NLR to be functional, we tested how many NLR-IDs from MIC1 and overall in the NLR phylogeny were paired with another NLR in head to head orientation (upstream NLR on the reverse strand, downstream NLR on the forward strand). Using our ‘tandem’ analysis workflow, we calculated the number of NLRs that had neighbouring NLRs. For each species, we compared results for tandem NLRs in head to head orientation from MIC1 to all NLRs.

Our results showed a significant enrichment (*p* < 0.05) of MIC1-NLRs being part of a NLR-pair in comparison to overall observations of paired NLRs (Table 3). This enrichment was most prominent in the two *Triticum* species, *T. aestivum* and *T. urartu*, followed by *S. italica*, *B. distachyon*, and *O. sativa*. Paired MIC1-NLRs in *A. tauschii* did not appear to be significantly enriched unless the search space was extended to include genes with insufficiently covered NB-ARC domains (see Methods – Phylogenetic Analysis). By contrast, we did not find any paired MIC-1 NLR in *H. vulgare* and *Z. mays*. While the results for the latter were likely due to the limited number of *Z. mays* NLRs in MIC1, the apparent lack of pairs in *H. vulgare* and lower numbers for *A. tauschii* were likely due to truncated assemblies where neighbouring genes might not be detected due to short scaffolds [25].

**Table 3.** Gene pair analyses.

| Pair search over all NLRs |  |  |  |  |  |
| --- | --- | --- | --- | --- | --- |
|  | MIC1 paired | MIC1 unpaired | Other paired | Other paired | *p* value |
| *A. tauschii* | 1 | 15 | 7 | 459 | 0.24 |
| *B. distachyon* | 5 | 7 | 29 | 214 | 0.013 |
| *H. vulgare* | 0 | 16 | 4 | 161 | 1.00 |
| *O. sativa* | 3 | 4 | 33 | 322 | 0.024 |
| *S. bicolor* | 2 | 4 | 20 | 236 | 0.082 |
| *S. italica* | 3 | 4 | 5 | 292 | 4.02e-04 |
| *T. aestivum* | 17 | 51 | 49 | 1616 | 9.26e-11 |
| *T. urartu* | 5 | 14 | 7 | 351 | 1.19e-04 |
| *Z. mays* | 0 | 0 | 0 | 84 | 1.00 |
| Pair search over all NLRs including short NB-ARC sequences |  |  |  |  |  |
|  | MIC1 paired | MIC1 unpaired | Other paired | Other paired | *p* value |
| *A. tauschii* | 4 | 12 | 24 | 723 | 0.0020 |
| *B. distachyon* | 8 | 4 | 41 | 344 | 1.06e-05 |
| *H. vulgare* | 0 | 16 | 7 | 242 | 1.00 |
| *O. sativa* | 5 | 2 | 61 | 471 | 4.13e-04 |
| *S. bicolor* | 2 | 4 | 29 | 329 | 0.085 |
| *S. italica* | 5 | 2 | 16 | 447 | 2.15e-06 |
| *T. aestivum* | 18 | 50 | 89 | 2671 | 1.61e-11 |
| *T. urartu* | 6 | 13 | 20 | 524 | 9.64e-05 |
| *Z. mays* | 0 | 0 | 6 | 174 | 1.00 |

When we mapped the location of pairs on the NLR phylogeny (Figure 3; Additional file 2), we observed that MIC1-NLRs paired exclusively with non-MIC1-NLRs, namely with members of clade C4 (35 pairs), clade C14 (1 pair), and the outer clade (1 pair). We further observed that the C4 and MIC1 clades were involved in more than half of all pairs (49, or respectively 37 out of 95). This was consistent with the pairing of RGA5 from MIC1 to RGA4 that is located in clade C4.

**Figure 3.**
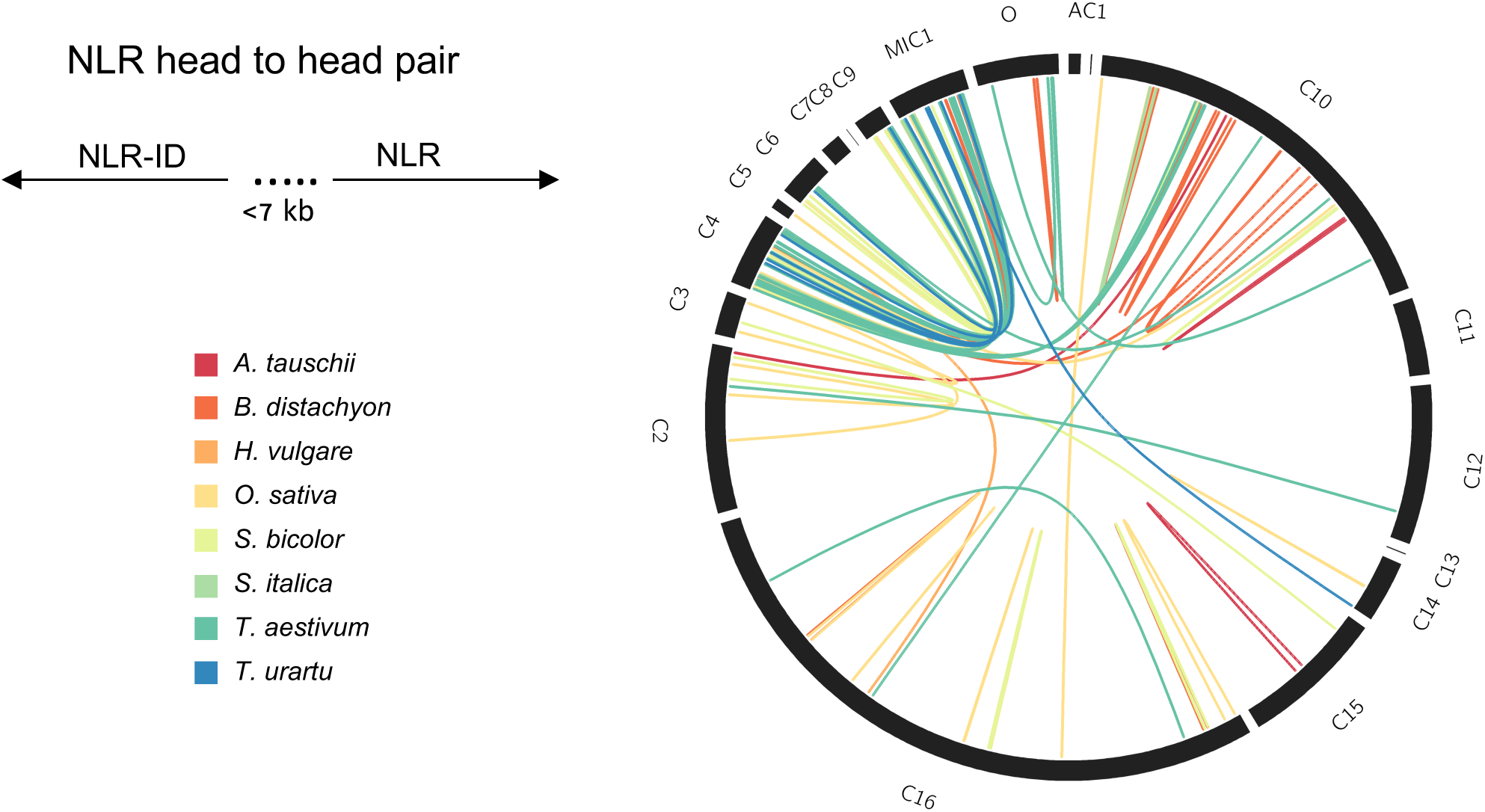
NLRs from MIC1 are genetically linked in head-to-head pairs with NLRs from clade 4 (C4) The schematic diagram (on the left) show the orientation and maximum distance of 7 kb that we used to identify NLR gene pairs. The Circos plot on the right shows links between NLRs from MIC1 and NLRs from other clades that are oriented across the plot in the same order as in the overall NLR phylogeny. NLR gene pair links from different species are color-coded as indicated in the legend on the left.

### Fusions associated with chromosomal rearrangement

We observed that NLRs from the MIC1 clade were found on different chromosomes across and within species. For five species analyzed in this study, the chromosomal location of NLR-IDs was available from the genome annotation. We tested for enrichment of NLR-IDs from the MIC1 clade on any particular chromosome and investigated whether these inter-species differences could be explained by whole-genome rearrangements during evolution (Additional File 3). Indeed, in most species NLR-IDs from the MIC1 clade were concentrated only on one or two chromosomes, such as chromosomes 2 and 5 in *S. bicolor*, chromosome 11 in *O. sativa*, chromosome 4 in *B. distachyon* and include 1:1 orthologs. These chromosomes form known syntenic blocks [23] indicating an ancient origin of the locus that was present in the common ancestor of grasses. However, none of the *O. sativa* or *B. distachyon* NLR-IDs in MIC1 shared conserved syntenic positions with NLR-IDs in the other species. We identified two examples in which NLR-IDs from *B. distachyon* are orthologous to non-fused NLRs in *O. sativa* (Figure 4A, Additional File 4). In both cases, the fusion events coincided with chromosomal rearrangements as was evident from the microsynteny analyses (Figure 4B, Additional File 4).

**Figure 4.**
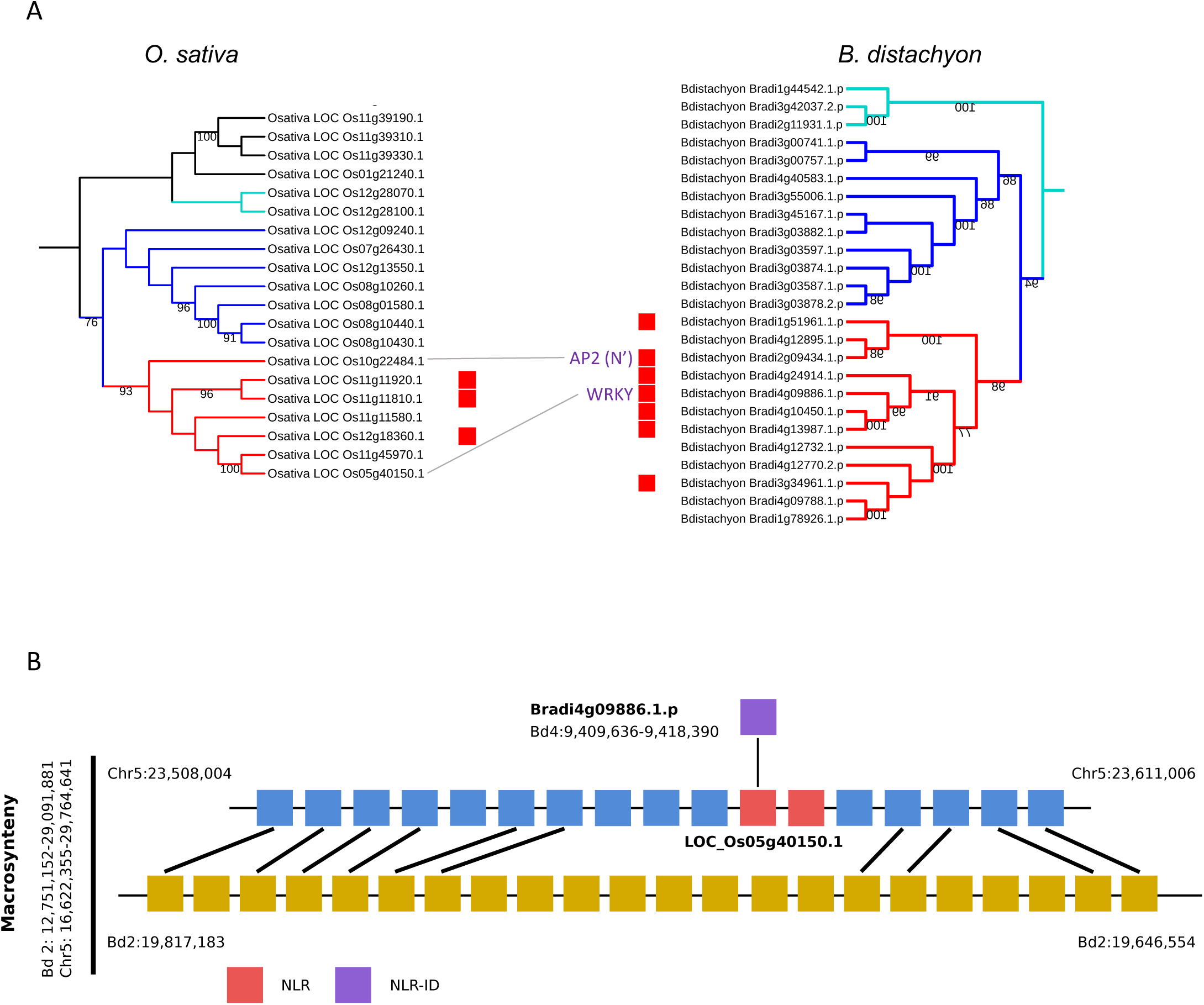
Orthologous NLRs from MIC1 in rice and Brachypodium show NLR gene coupled to generation of new NLR-IDs. (A) Phylogeny of NLRs from MIC1 in rice and Brachypodium based on the NB-ARC domain. Red boxes indicate NLRs with IDs. The links between trees highlight orthologous NLRs between rice and Brachypodium (1:1 orthologs). (B) Microsynteny analyses between rice and Brachypodium. Blue boxes and yellow ochre boxes represent syntenic genes in rice and Brachypodium, respectively. Red boxes indicate NLRs, purple boxes – NLR-IDs.

With the proliferation of the NLR-ID MIC1 clade in wheat, we also observed more divergent locations of NLR-IDs, mostly concentrated on chromosomes 7AS, 7DS, and 4AL, but also conserved across all three subgenomes on the homoeologous chromosome groups 1, 3 and 6 (Additional File 5). Such proliferation can be explained by more recent large scale genomic rearrangements, such as the translocation of a chromosomal region from 7BS to 4AL and other known chromosomal translocation and duplication events [26, 27]. The expansion of the MIC1 clade in the *Triticeae*, its proliferation to multiple genomic locations as well as the increased diversity of integrated domains might be linked to the overall increased fraction of TEs in these genomes compared to smaller genomes of other grasses.

### Frequent domain swapping at orthologous NLRs

To further understand the evolution of ID fusions, we aligned the proteins from the MIC1 clade and the associated outer and ancestral clades and reconstructed the phylogeny of these clades alone by a maximum likelihood approach (Figure 5; Additional File 6). Each gene was annotated with a figure showing the positions of canonical (NB and LRR) and non-canonical domains. One of the first striking observations from this representation were the differences in the distribution and diversity of the ID domain(s) amongst the proteins in the ancestral (cyan), outer (blue) and inner MIC1 (red) clade. The ancestral clade had no ID domains, the outer clade had three groups of proteins (all from wheat or its progenitors) with ID domains at their N-terminal ends. In each of these groups, the ID domains primarily originated from only three protein domain families, WRKY and Motile_Sperm, which in one group can be linked to a Pkinase(_Tyr) domain. The presence of the same domain(s) in each group has parallels with proteins from other clades of the NB-ARC family – for example clades 2 and 3 in Figure 1A - that have an identical integrated domain from one family, suggesting that there has been a single integration event followed by gene duplication. There are several examples, notably in the *Triticeae*, in which genes, including homoeologs of the same gene triad, share the same domain at the same position in the protein. Among such examples are NLR-GRAS, NLR-kinase and NLR-NPR1 (Figure 5). This conservation in architecture indicates a common ancestry and selection to maintain a functional fusion. In contrast to these patterns, the majority of closely related NLR proteins within the inner clade have diverse ID domains, where domains are derived from different protein families and primarily exist near the C-terminal end of the NLR. In some closely related genes, a different domain resides in a similar position indicating that there is a common integration point in these genes.

**Figure 5.**
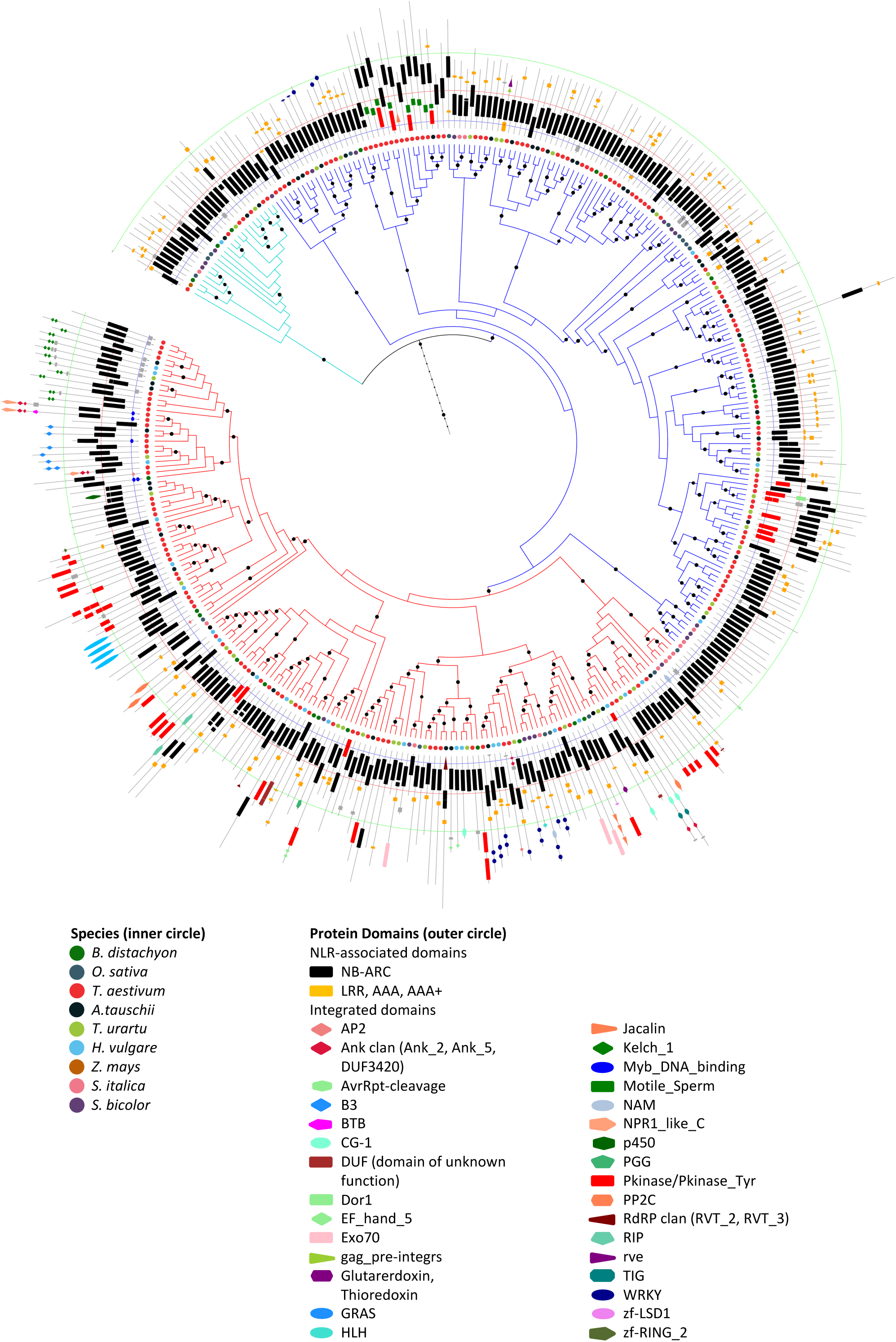
Close-up of MIC1 displaying rapid domain recycling. The branches of the hotspot clade, the outer clade and the ancestral clade are shown in red, blue and cyan respectively. Dots on the branches indicate a bootstrap support value ≥ 85 %. Alongside the tree are cartoons of each protein, annotated with the domain(s) in the position that they appear in the protein (protein backbone, grey line; NB-ARC domain, black rectangle; LRR and AAA, TIR and RPW8 domains, orange rectangles; other domains in different colours and shapes as indicated in the key). E-value cut-off for presence of an ID domain, 0.001; domains with e-value > 0.001 and ≤ 0.05 are shown as grey rectangles. E-value cut-off for an LRR, AAA, TIR or RPW8 domain, 10.0.

Such domain swapping raised questions about the mechanisms by which domain integration can be achieved and maintained. For example, did these proteins share sequences that increase the likelihood of integration events and are there NLR/ID combinations that create a functionally successful NLR protein that becomes fixed in the species?

### MIC1-NLR-IDs share a protein motif at the site of domain integrations

In order to look for shared sequences that might answer the questions above, we searched for protein motifs that were enriched in the MIC1 clade. For every protein, we extracted all regions without a domain annotation from InterProScan. Motif prediction using MEME found seven motifs (I06, I07, I08, I09, I11, I17, and I40) that were saturated within the MIC1 clade (Additional File 7). Two motifs, I09 and I11 were associated with the region between the coiled-coil (CC) and NB domains, whereas I07, I40, I17, I08, and I06 were associated with the LRR region and/or between the LRR and ID (Figure 6A, C-terminal ID). Motif I09 was widespread, with 80 of 159 proteins in the MIC1 clade harboring this motif, whereas motif I11 occured in 63 proteins in the MIC1 clade. When found together, they were always in tandem (I09-I11) and were located upstream of the NB domain. Motifs I09 and I11 could be regions of the subfamily of CCs and/or NBs found in proteins within the MIC1 clade that were not annotated by InterProScan.

**Figure 6.**
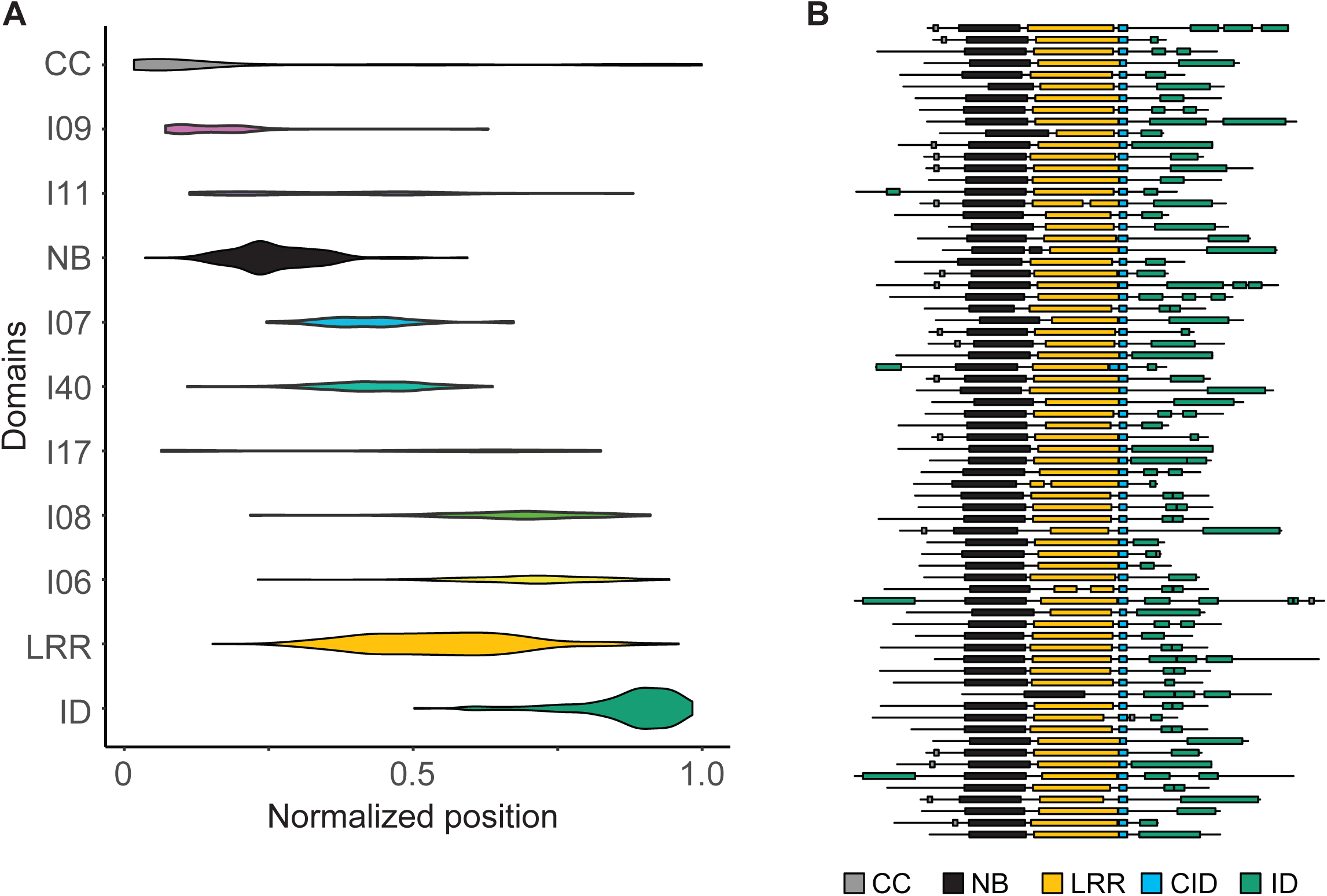
NLR-IDs from MIC1 share a protein motif at the site of domain integrations. (A) Distribution of CC, NB, LRR, and ID domains and motifs identified using MEME on unannotated regions of NLRs within the MIC1 clade with C-terminal ID. For every NLR, the length of the NLR was normalized to 1.0 and the midpoint of identified domains was normalized to protein length. (B) Domain structure of 70 NLR-IDs within the MIC1 clade that contain the CID domain. The CID domain is located immediately upstream of the site of integration.

For the group of motifs located between the NB and ID domains, we found that I07, I40, I17, I08 were LRR motifs trained on regions that were not annotated based on InterProScan analysis. These could therefore be excluded from further analysis. In contrast, I06 was a motif specifically associated with NLRs in the MIC1 clade and was located immediately upstream of the integrated domains. Based on its conservation and association with IDs, we designated this domain the CID domain. We developed a Hidden Markov Model trained on the CID domain (Additional files 8 and 9) and overlaid its presence/absence on the phylogenetic tree of NLRs (Additional file 10). The CID domain was present in the majority of genes (70%) within the MIC1 clade, occasionally found (20%) in genes in the outer MIC1 clade, and found in nine genes outside these clades. Alignment of the regions encompassing the CID and ID domains uncovered a clear breakpoint between these domains for the majority of NLRs in the MIC clade (Figure 6B). This suggests that while different domains may integrate within NLRs in the MIC1 clade, selection acts to maintain the CID domain prior to the integration site of IDs.

### Duplication of genes encoding IDs followed by translocation of either ID or NLR lead to new NLR-ID formation

Translocation of NLRs in the comparison between rice and *B. distachyon* provided an initial understanding of the origin of new NLR-IDs. To further elucidate the mechanisms of NLR-ID formation, we looked for examples of the most recent integration. A polyploid bread wheat (*T. aestivum*), presented an ideal system for these analyses. Wheat has an elevated number of NLRs and NLR-IDs (Figure 2A), a high incidence of new integrations as well as multiple orthologous copies of each gene (A, B, and D). The presence of homoeologs allowed us to trace the origin of the ID as well as the translocation of NLR or ID.

To identify proteins that were most closely related to the donor ID domains in *T. aestivum* NLR-IDs, we constructed phylogenies for eight families that harbor ID domains: AP2/ERF, Exo70, GRAS, Kelch, NPR1_like_C, Pkinase, Pkinase_Tyr and WRKY (Additional File 11, Additional File 12). We uncovered multiple independent integrations of donor proteins into different NLR proteins, particularly for WRKY and Pkinase. The majority of domains existed as complete domains within the NLR protein suggesting that they might have retained their orginal function and might provide access to existing signaling networks; two transcription factor families, AP2/ERF and WRKY included ID donor proteins that are already known to be involved in stress and pathogen response [28].

We considered three possible mechanisms of NLR-ID formation (1) retrotransposition of cDNA derived from the parental gene, (2) transposition of the parental gene, and (3) ectopic recombination during which double-stranded DNA breaks are repaired using a non-homologous exogenous parental gene as a template. All three mechanisms have been observed previously in cereal genomes [29], and both retrotransposition and ectopic recombination have been suggested as diversification mechanisms of NLRs [30]. We extracted the coding DNA sequences of integrated domains for 40 *T. aestivum* NLR-ID genes from MIC1 and aligned them back to the genome (BLASTN, e-value 1e-3). Similar to the NLR portion of the genes, the majority of integrated domains contained introns. Therefore, we conclude that retrotransposition of IDs is unlikely.

We looked at a recent exchange of IDs in NLR-IDs, specifically at NLR-WRKY/AP2a (Figure 7) and MYB/AP2b-NLR domain swaps (Additional File 13), to further understand how exogenous domains become fused to NLRs. In case of NLR-WRKY/AP2a swap, there were three homoeologous NLRs on chromosomes 5A, 5B, and 5D, respectively with distinct C-terminal fusions (Figure 7A). The AP2 domain in the NLR-ID on chromosome 5BL replaced a more ancient WRKY domain integration present in other wheat homoeologs on 5AL and 5DL as well as in other grasses (Figure 7A). Therefore, the integration of AP2a occurred after the split of the diploid wheat genome progenitors (<2.7 MYA) [31]. The closest homolog of the AP2 ID was located on chromosome 3 and was present in all subgenomes (A, B and D), suggesting a duplication of the parental ID copy either before or coupled with movement into the NLR (Figure 7B). By aligning the AP2a nucleotide sequence to its parental genes on chromosome 3 with BLASTN, we observed that integration involved a part of AP2 intron 1 and exon 2, which became fused with the intron 2 of NLR displacing the WRKY gene (Figure 7C). Since the three parental AP2 genes were intact and there are no additional paralogs on any sub-genome, we concluded that the gene must have been copied first into a new location if transposition had been involved. We found no evidence of the residual first exon of AP2 on any wheat subgenome. A recent transposition would also have left a footprint, such as terminal inverted repeats. A BLASTN search of the ID sequence and its surrounding region against itself revealed only a very short terminal inverted repeat (TATAGCTACAG) on each side of ID. The presence of short TIRs suggests that if transposition had been involved, it would have been mediated through a poorly characterized class of DNA transposon, such as hAT or PIF/Harbinger. The linker region between the NB-ARC and AP2 domains contains an 829 bp region, which lacks homoeologs in the wheat genome and contains the 86 bp inverted repeat.

**Figure 7.**
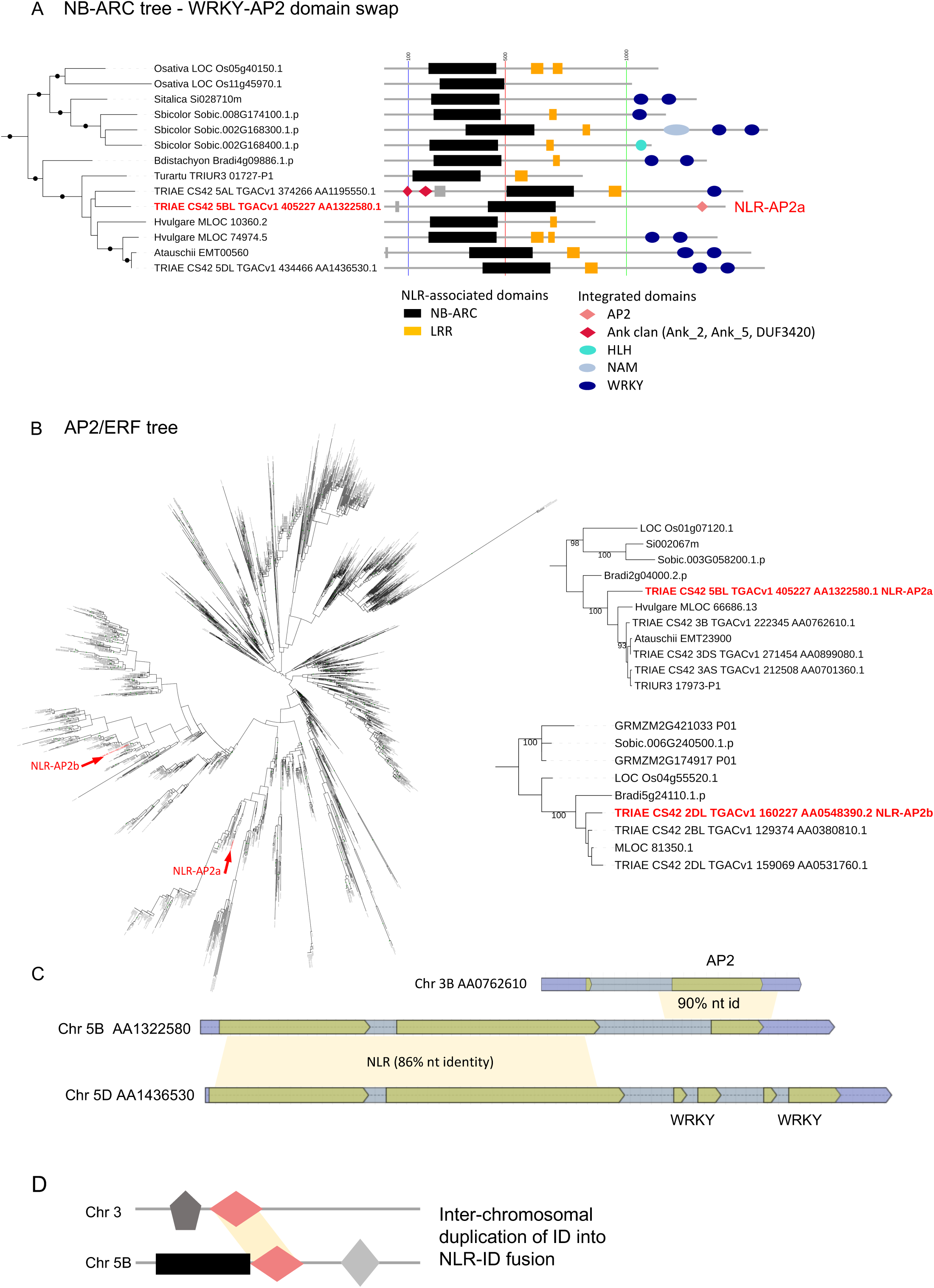
NLR-WRKY/AP2a domain swap involved inter-chromosomal copy-and-paste of the AP2 gene. (A) A clade in the phylogeny of NLR-ID proteins from Figure 5 that includes wheat A, B and D genome homoeologs from the same genetic position. A swap is evident between an AP2 and a WRKY domain (highlighted by red boxes). Dots on the tree branches indicate a bootstrap support value ≥85 %. E-value cut-off for presence of an ID domain, 0.001; a domain with e-value > 0.001 and ≤0.05 is shown as a grey rectangle. E-value cut-off for an LRR domain, 10.0. (B) An AP2/ERF family tree (left) showing two clades that contain an NLR acceptor ID protein (NLR-AP2a and NLR-AP2b, indicated in red). The protein sequences of these clades were re-aligned and the trees re-estimated (right) to confirm the identity of the donor protein, evident from the high bootstrap support values. (C) Protein alignment cartoon of one of the AP2 donor proteins and the acceptor protein, NLR-AP2a. By contrast, the NLR acceptor protein homoeologs contain two WRKY domains in their C-terminal ends.

In the second example, an AP2 domain displaced the MYB domain as N-terminal fusion of NLR (Additional File 13). The AP2b gene was evolutionary distinct from AP2a (Figure 7B), representing an independent fusion of a distant family member. Interestingly, in this example, it was the NLR that moved into new location since the AP2b-NLR is located on chromosome 2DL and its NLR homoeologs were located on chromosome 7AS and, via a known large-scale chromosomal translocation from 7BS [32], on chromosome 4AL (Additional file 13). Moreover, chromosome 2DL contained a non-fused copy of AP2b gene indicating that, as in the case of AP2a, the parental ID gene was duplicated before integration. While we saw more examples of distinct IDs fused to orthologous NLRs (Figure 6), this wheat example together with our rice/*B. distachyon* analyses suggested that either the ID or NLR can be translocated to a distinct genomic location to create a new fusion event.

Altogether, the data suggest that the integration of exogenous domains into NLRs follows a gene duplication plus inter-chromosomal gene translocation mechanism. The most likely mechanism of such ‘copy-and-paste’ is ectopic recombination, although other means involving copy number increase and DNA transposition cannot be excluded.

### Discussion

We have investigated the formation of NLR-IDs in grasses and demonstrated that while many NLR clades are capable of new domain integrations, the distribution of NLR-IDs is uneven across the NLR phylogeny. While some clades rich in NLR-IDs represent the proliferation of single ancient domain integrations, one dominant clade MIC1 harbors the most diverse NLR fusions. MIC1 includes several known NLR-IDs, rice RGA5 and Pi-ta, as well as barley Rpg5. The NLRs in MIC1 are often genetically linked to another NLR originating from a distinct clade, C4, which includes the *RGA5* gene partner, *RGA4*. In MIC1, new NLR formation is an active mechanism that involves inter-chromosomal gene movement. Our synteny analyses between rice and *B. distachyon* documented movement of NLRs that lead to acquisition of their IDs. Our analyses in wheat showed the occurrence of domain swaps in NLR-IDs. Although we cannot exclude transposition as a mechanism for NLR-ID formation, we have not observed any known TE or TE-associated inverted repeats linked to gene transposition. Therefore, our data suggest that ectopic recombination is the most likely mechanism of inter-chromosomal gene transfer.

How the innate immune system of plants is capable of acquiring new pathogen recognition specificities remains a key question in our understanding of plant-pathogen interactions. Its answer is closely linked to the evolution of plant immune receptors and their diversification. The processes underpinning genome evolution often include domain duplication, fission and fusion [33], which have recently been implicated in NLR evolution [7-9, 34]. One of the most advantageous pathways to recognize multiple pathogens is by guarding common plant proteins that are targeted by multiple, if not all, pathogens. However, a mechanism that involves self-recognition can quickly lead to auto-immunity [35, 36]. Genetic linkage of NLRs and the proteins they guard into NLR-IDs can prevent allele shuffling and autoimmunity as well as enable coordinated transcriptional and translational regulation that is important for controlling protein stoichiometry and co-evolution. Similarly, genetic linkage of an NLR and its helper in a gene pair can be beneficial. Therefore, a clade of NLRs that gained the capacity to ‘integrate’ new proteins – either by allowing higher levels of DNA exchange or being able to accept different fusions without autoactivity, or the combination of both – presents an evolutionary advantage.

Our model (Figure 8) summarizes how these processes could have driven the expansion and diversification during evolution of the proteins in NLR-ID MIC1. While the ability to be amenable to new integration likely preceded the origin of grasses, the expansion of the MIC1 in grass genomes enabled diverse fusions and rapid sampling of new fusions. Subsequent to the ancestral genes’ tolerance and propensity to form fusions, perhaps through the evolution of novel domains such as the CID protein motif, diversification occurred, resulting in the large variety of exogenous integrated domains.

**Figure 8.**
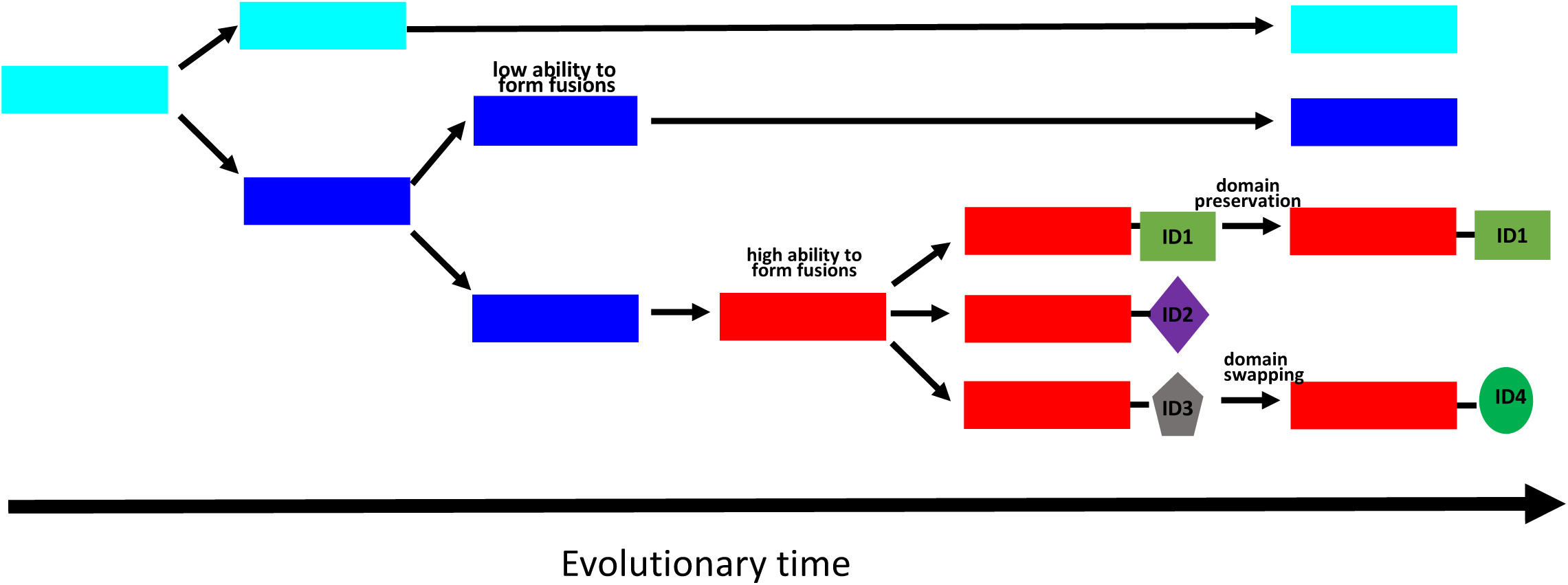
Evolutionary model of NLR-ID hotspot formation and diversification. Arrows indicate NLR fate over evolutionary time; orange arrows indicate duplication events. The ancestral protein (cyan) underwent duplication to form the outer clade of proteins (blue). Then proteins in specific clades, especially those proteins in hotspot 1 (red) gained the ability to form fusions with other domains (ID’s). Some proteins have maintained the same domain (for example, ID1) but other proteins have undergone further diversification through the exchange of the ID domain (for example, ID3 and ID4).

In the future, the availability of higher quality genome assemblies as well as multiple genomes for each species will allow more detailed analyses of syntenic gene clusters and will identify the precise locations of DNA breakpoints that lead to NLR-ID formation. Combining long molecule sequencing RenSeq [37] with population genetics analyses will allow us to estimate how rapidly new gene fusions are formed within populations and how fast the selection of advantageous combinations occurs in nature.

There is an urgent need for new genetic sources of resistance for future sustainable crop production [38, 39]. Our identification of NLRs that are highly amenable to the integration of exogenous domains can be efficiently exploited for advancing the understanding of how new immune receptor specificities are formed and provide new avenues to generate novel synthetic fusions.

## Methods

### Identification of NLRs and NLR-IDs in plant genomes

NLR plant immune receptors were identified in nine monocot species by the presence of the common NB-ARC domain (Pfam PF00931) as described previously [9]. The *T. aestivum* (TGAC v1) and *A. tauschii* genomes (ASM34733v1) were downloaded from EnsemblPlants and analyzed using the same pipeline used previously [9]. This analysis included the identification of proteins with ‘integrated’ domains (Additional File 14) All scripts are available from <u>https://github.com/krasileva-group/plant_rgenes</u>, script versions used in this study include K-parse_Pfam_domains_NLR-fusions-v2.4.pl and K-parse_Pfam_domains_v3.1.pl

### Phylogenetic Analysis

An HMM - based on the Pfam model PF00931 - was built to include the ARC2 subdomain which is also present in plant NB-ARC proteins (Additional File 15 and 16). To build the model of NB-ARC1-ARC2, eight proteins (SwissProt identifiers: APAF_HUMAN, LOV1A_ARATH, K4BY49_SOLLC, RPM1_ARATH, R13L4_ARATH, RPS2_ARATH, DRL24_ARATH, DRL15_ARATH) were aligned using PRANK [40] and the HMM was built from this alignment with HMMER3 HMMBUILD [41], using default parameters for both programs. Amino acid sequences encoding the NB-ARC proteins were aligned to this HMM using the HMMER3 HMMALIGN program (version 3.1b2) [41]. The resulting alignment of the NB-ARC1-ARC2 domain was converted to fasta format using the HMMER ESL-REFORMAT program. Any gap columns in the alignment of target proteins with the HMM were removed. Sequences with less than 70% coverage across the alignment were removed from the data set to reduce false placement in the tree of sequences with insufficient coverage across the domain. The longest sequence for each gene out of the available set of splice versions was used for phylogenetic analysis. In addition, 38 proteins encoding genes with characterized and known functions in pathogen defence from the literature were also included; the list of genes was based on a curated R-gene dataset by Sanseverino et al, 2012 (<u>http://prgdb.crg.eu</u>). Phylogenetic analysis was carried out using the MPI version of the RAxML (v8.2.9) program [42] with the following method parameters set: -f a, -x 12345, -p 12345, -# 100, -m PROTCATJTT. The tree contained 4,133 sequences, 338 columns, took 67 hours to generate and required 17 GB RAM. Separate trees for each species were also prepared using the same methods (Additional File 1). The overall species phylogeny was constructed using NCBI taxon identification numbers at phyloT (phylot.biobyte.de). The trees were mid-point rooted and visualized using the Interactive Tree of Life (iToL) tool [43] and are publicly available at <u>http://itol.embl.de</u> under ‘Sharing data’ and ‘KrasilevaGroup’ and in Newick format in Additional File 17. Annotation files were prepared for displaying the presence of ID domains in the proteins, identifying species gene identifiers by colour and visualizing the location of individual domains within the protein backbone. An ID domain was defined as being any domain, except for LRR, AAA, TIR, and RPW8, which are often associated with NB-ARC-containing proteins.

To identify donor genes for the ID domains of the NLR-ID proteins, phylogenetic trees for eight donor gene families (AP2, Exo70, GRAS, Kelch, NPR1_like_C, Pkinase, Pkinase_Tyr and WRKY) were produced by the methods described above, except that the species chosen were *A. thaliana*, *M. truncatula*, *B. distachyon*, and *T. aestivum*, except for the ERF family for which *A. thaliana*, *M. truncatula*, and all nine monocot species described above for the NLR family were included. The following protein annotation files were used: *A. thaliana (*TAIR10_pep_20101214_updated (TAIR10)), *B. distachyon* (Bdistachyon_314_v3.1.protein.fa (Phytozome, version 12)), *M. truncatula (*Mtruncatula_285_Mt4.0v1.protein.fa (Phytozome, version 12)), and *T. aestivum* (TGAC_v1 protein annotation as described above (EnsemblPlants website)), respectively. The HMMs used for each family were taken from the Pfam-A database (Release30), except for the model for the AP2/ERF domain, which was created from an alignment with PRANK of *A. thaliana* and rice ERF proteins.

### Identification of protein motifs

NLRs within the MIC1 clade were annotated for known domains using InterProScan (v5.20-59.0). Domains were annotated and undefined regions within NLRs were extracted using the QKdomain pipeline (<u>https://github.com/matthewmoscou/QKdomain)</u>. All undefined regions were required to be at least 20 amino acids long. MEME (v4.11.2) was used for motif prediction using the undefined regions [44]. FIMO was used to identify motifs in the entire set of NLRs from diverse grass species [44]. Visualization of the presence/absence of motifs was performed using iTOL [43]. Multiple sequence alignments were performed using MUSCLE (v3.8.31) [45]. HMMER3 (v3.1b1) HMMBUILD was used to train Hidden Markov Models on conserved sequences and HMMSEARCH was used to search the entire NLR dataset, using default parameters [41]. The complete pipeline, including scripts and data sets, is available from the Github repository NLR-ID_motif (<u>https://github.com/matthewmoscou/NLR-ID_motif)</u>.

### Detection of Paired NLRs

Gene annotations of eight grass species (Aegilops_tauschii.ASM34733v1.33.gff3, Bdistachyon_283_v2.1.gene.gff3, Hordeum_vulgare.ASM32608v1.33.gff3, Osativa_204_v7.0.gene.gff3, Sbicolor_255_v2.1.gene.gff3, Sitalica_164_v2.1.gene.gff3, and Triticum_urartu.ASM34745v1.33.gff3, Zmays_284_6a.gene.gff3) were obtained from Phytozome V11. The gene annotation for *T. aestivum* was obtained from <u>http://opendata.earlham.ac.uk/Triticum_aestivum/TGAC/v1/annotation/Triticum_aestivum_CS4</u> 2_TGACv1_scaffold.annotation.gff3.gz [24].

NLR genes were tested for the presence of paired NLRs (NLR1 upstream of NLR2, NLR1 in reverse orientation, NLR2 in forward orientation, and a maximum distance of 15 kbp between NLRs). The paired NLR search was based on bedtools closest (with -d and -k2 parameters) [46], implemented in a script called tandem (<u>https://github.com/krasileva-group/tandem)</u> and the results were displayed using Circos [47]. Statistical significance was calculated with Fischer’s exact test (as implemented in scipy) by comparing the number of genes involved in NLR tandems in the MIC1 clade with the set of all NLRs.

## Additional Files

**Additional File 1**: Maximum likelihood phylogeny based on the NB-ARC domain of all NLRs and NLR-IDs for each of the nine grass species under study. (A) *S. italica* (B) *S. bicolor* (C*) Z. mays* (D) *O. sativa* (E) *B.distachyon* (F) *H.vulgare* (G) *A.tauschii* (H) *T. aestivum* (I) *T.urartu*. Proteins with integrated domains are represented by red squares. Branch colour, arc colour and labels represent the MIC1 clade as defined in Figure 1. (.pptx)

**Additional File 2:** Clade memberships of all NLRs present in Figure 1 tree. (.tsv)

**Additional File 3**: Chromosome distributions of the NLRs identified from five monocot species. (.xlsx)

**Additional File 4: Orthologous NLRs from MIC1 in rice and *Brachypodium distachyon*** (A) A Circos plot summarizing the locations of ortholgous NLRs between rice and *B. distachyon* (B) Microsynteny analyses between rice and *B. distachyon*. Blue and yellow ochre boxes represent syntenic genes in rice and *B. distachyon*, respectively. Red boxes indicate NLRs, purple boxes indicate NLR-IDs. (.pdf)

**Additional File 5**: Genetic locations of all NLRs and NLR-IDs present in the tree in Figure 1A, anchored to the genetic map of *T. aestivum* CS42. (.tsv)

**Additional File 6**: Gene identification numbers for NLRs from the MIC1 clade, blue neighbouring clade, and cyan ancestral clade. (.txt)

**Additional File 7:** Motifs identified using MEME that are associated with MIC1 clade. (.pdf)

**Additional File 8:** Alignment file used to generate HMM for CID domain. (.txt)

**Additional File 9:** The HMM for the CID domain. (.txt)

**Additional File 10:** Presence/absence of motifs (marked as black dots) relative to the NB phylogenetic tree. (A) I06, (B) I09, and (C) I11. (.pdf)

**Additional File 11**: Maximum likelihood phylogeny for eight gene families containing proteins with ID domains that were used to identify potential donor genes within each family for *T. aestivum* NLR-ID genes from hotspot clade 3. The gene identifiers highlighted in red are the acceptor genes containing an NB-ARC domain. (A) AP2/ERF family (B) Exo70 family (C) GRAS family (D) Kelch family (E) NPR1_like_C (F) Pkinase (G) Pkinase_Tyr (H) WRKY (.pptx)

**Additional File 12:** Potential donor - NLR acceptor gene sets, as observed from phylogenetic trees of the donor ID genes. High bootstrap support was used to determine likely donor ID and acceptor NLR-ID gene clades. (.xlsx)

**Additional File 13:** The AP2b/MYB-NLR domain swap includes duplication of AP2 gene and inter-chromosomal transfer of NLR. (.pptx)

**Additional File 14**: Integrated domains found in the NLR proteins of nine grass species at relaxed e-value cutoff (<0.05). (.txt)

**Additional File 15**: The alignment of NB, ARC1, and ARC2 used to train the HMM for NLR proteins in phylogenetic analysis. (.txt)

**Additional File 16**: The HMM trained from alignment of NB, ARC1, and ARC2 for NLR proteins in phylogenetic analysis. (.txt)

**Additional File 17**: All NLR phylogenies in Newick format. (.txt)

## Declarations

### Ethics approval and consent to participate

Not applicable

### Consent for publication

Not applicable

### Availability of data and materials

The datasets supporting the conclusions of this article are included within the article and its additional files.

## Authors contributions

PB, WH, MM and KVK designed the study. PB performed phylogenetic analyses, GD collected a set of well-studied NLRs, EB and KVK performed domain analyses, CS, PB and KVK performed ‘tandem’ analyses, CS and WH performed synteny analyses, WJ and MM analyzed protein motifs, PB and KVK analyzed NLR-AP2 fusions. All authors contributed to writing of the manuscript.

## Acknowledgements

Authors are grateful to all members of Krasileva group and many colleagues, especially Sophien Kamoun, for thoughtful discussions of the presented material. We thank Daniil Prigozhin for suggestions on data analyses and the manuscript. The high-performance computing resources and services used in this work were supported by the EI Scientific Computing group alongside the NBIP Computing infrastructure for Science (CiS) group.

## References

1. Jones JDG, Vance RE, Dangl JL: Intracellular innate immune surveillance devices in plants and animals. Science 2016, 354:aaf6395–aaf6395.

2. Dodds PN, Rathjen JP: Plant immunity: towards an integrated view of plant– pathogen interactions. Nat Rev Genet 2010, 11:539–548.

3. Hall Sa, Allen RL, Baumber RE, Baxter La, Fisher K, Bittner-Eddy PD, Rose LE, Holub EB, Beynon JL: Maintenance of genetic variation in plants and pathogens involves complex networks of gene-for-gene interactions. Mol Plant Pathol 2009, 10:449–457.

4. Joshi RK, Nayak S: Perspectives of genomic diversification and molecular recombination towards R-gene evolution in plants. Physiol Mol Biol Plants 2013, 19:1–9.

5. Le Roux C, Huet G, Jauneau A, Camborde L, Trémousaygue D, Kraut A, Zhou B, Levaillant M, Adachi H, Yoshioka H, et al: A receptor pair with an integrated decoy converts pathogen disabling of transcription factors to immunity. Cell 2015, 161:1074–1088.

6. Sarris PF, Duxbury Z, Huh SU, Ma Y, Segonzac C, Sklenar J, Derbyshire P, Cevik V, Rallapalli G, Saucet SB, et al: A plant immune receptor detects pathogen effectors that target WRKY transcription factors. Cell 2015, 161:1089–1100.

7. Cesari S, Bernoux M, Moncuquet P, Kroj T, Dodds PN: A novel conserved mechanism for plant NLR protein pairs: the “integrated decoy” hypothesis. Front Plant Sci 2014, 5:606.

8. Kroj T, Chanclud E, Michel-Romiti C, Grand X, Morel JB: Integration of decoy domains derived from protein targets of pathogen effectors into plant immune receptors is widespread. New Phytol 2016, 210:618–626.

9. Sarris PF, Cevik V, Dagdas G, Jones JDG, Krasileva KV: Comparative analysis of plant immune receptor architectures uncovers host proteins likely targeted by pathogens. BMC Biol 2016, 14:8.

10. Césari S, Kanzaki H, Fujiwara T, Bernoux M, Chalvon V, Kawano Y, Shimamoto K, Dodds P, Terauchi R, Kroj T: The NB-LRR proteins RGA4 and RGA5 interact functionally and physically to confer disease resistance. EMBO J 2014, 33:1941 LP –1959.

11. Saucet SB, Ma Y, Sarris PF, Furzer OJ, Sohn KH, Jones JD: Two linked pairs of Arabidopsis TNL resistance genes independently confer recognition of bacterial effector AvrRps4. Nat Commun 2015, 6:6338.

12. Prasad V, Stromberg C, Alimohammadian H, Sahni A: Dinosaur coprolites and the early evolution of grasses and grazers. Science 2005, 310:1177–1180.

13. Prasad V, Strömberg CaE, Leaché aD, Samant B, Patnaik R, Tang L, Mohabey DM, Ge S, Sahni a: Late Cretaceous origin of the rice tribe provides evidence for early diversification in Poaceae. Nat Commun 2011, 2:480.

14. Dubcovsky J, Dvorak J: Genome plasticity a key factor. Science 2007, 316:1862–1866.

15. Moore G, Devos KM, Wang Z, Gale MD: Cereal genome evolution. Grasses, line up and form a circle. Curr Biol 1995, 5:737–739.

16. Zhang Y, Xia R, Kuang H, Meyers BC: The diversification of plant NBS-LRR defense genes directs the evolution of MicroRNAs that target them. Mol Biol Evol 2016, 33:2692–2705.

17. Muñoz-Amatriaín M, Eichten SR, Wicker T, Richmond TA, Mascher M, Steuernagel B, Scholz U, Ariyadasa R, Spannagl M, Nussbaumer T, et al: Distribution, functional impact, and origin mechanisms of copy number variation in the barley genome. Genome Biol 2013, 14:R58.

18. Periyannan SK, Moore J, Ayliffe M, Bansal U, Wang X, Huang L, Deal K, Luo M, Kong X, Bariana H, et al: The gene Sr33, an ortholog of barley Mla genes, encodes resistance to wheat stem rust race Ug99. Science 2013, 341:786–788.

19. Cesari S, Thilliez G, Ribot C, Chalvon V, Michel C, Jauneau A, Rivas S, Alaux L, Kanzaki H, Okuyama Y, et al: The rice resistance protein pair RGA4/RGA5 recognizes the Magnaporthe oryzae effectors AVR-Pia and AVR1-CO39 by direct binding. Plant Cell 2013, 25:1463–1481.

20. Brueggeman R, Druka A, Nirmala J, Cavileer T, Drader T, Rostoks N, Mirlohi A, Bennypaul H, Gill U, Kudrna D, et al: The stem rust resistance gene *Rpg5* encodes a protein with nucleotide-binding-site, leucine-rich, and protein kinase domains. Proc Natl Acad Sci USA 2008, 105:14970–14975.

21. Bryan GT, Wu KS, Farrall L, Jia Y, Hershey HP, McAdams SA, Faulk KN, Donaldson GK, Tarchini R, Valent B: A single amino acid difference distinguishes resistant and susceptible alleles of the rice blast resistance gene Pi-ta. Plant Cell 2000, 12:2033–2046.

22. Costanzo S, Jia Y: Alternatively spliced transcripts of Pi-ta blast resistance gene in Oryza sativa. Plant Sci 2009, 177:468–478.

23. Vogel JP, Garvin DF, Mockler TC, Schmutz J, Rokhsar D, Bevan MW, Barry K, Lucas S, Harmon-Smith M, Lail K, et al: Genome sequencing and analysis of the model grass Brachypodium distachyon. Nature 2010, 463:763–768.

24. Clavijo BJ, Venturini L, Schudoma C, Accinelli GG, Kaithakottil G, Wright J, Borrill P, Kettleborough G, Heavens D, Chapman H, et al: An improved assembly and annotation of the allohexaploid wheat genome identifies complete families of agronomic genes and provides genomic evidence for chromosomal translocations. Genome Res 2017, 27:885–896.

25. International Barley Genome Sequencing Consortium: A physical, genetic and functional sequence assembly of the barley genome. Nature 2012, 491:711–716.

26. Salse J, Bolot S, Throude M, Jouffe V, Piegu B, Quraishi UM, Calcagno T, Cooke R, Delseny M, Feuillet C: Identification and characterization of shared duplications between rice and wheat provide new insight into grass genome evolution. Plant Cell 2008, 20:11–24.

27. Clavijo BJ, Venturini L, Schudoma C, Garcia Accinelli G, Kaithakottil G, Wright J, Borrill P, Kettleborough G, Heavens D, Chapman H, et al: An improved assembly and annotation of the allohexaploid wheat genome identifies complete families of agronomic genes and provides genomic evidence for chromosomal translocations. bioRxiv 2016: http://biorxiv.org/content/early/2016/2010/2013/080796.

28. Eulgem T, Rushton PJ, Robatzek S, Somssich IE: The WRKY superfamily of plant transcription factors. Trends Plant Sci 2000, 5:199–206.

29. Wicker T, Buchmann JP, Keller B: Patching gaps in plant genomes results in gene movement and erosion of colinearity. Genome Res 2010, 20:1229–1237.

30. Leister D, Kurth J, Laurie Da, Yano M, Sasaki T, Devos K, Graner a, Schulze-Lefert P: Rapid reorganization of resistance gene homologues in cereal genomes. Proc Natl Acad Sci USA 1998, 95:370–375.

31. Dvorak J, Akhunov ED: Tempos of gene locus deletions and duplications and their relationship to recombination rate during diploid and polyploid evolution in the Aegilops-Triticum alliance. Genetics 2005, 171:323–332.

32. Devos KM, Dubcovsky J, Dvorak J, Chinoy CN, Gale MD: Structural evolution of wheat chromosomes 4A, 5A, and 7B and its impact on recombination. Theor Appl Genet 1995, 91:282–288.

33. Moore AD, Björklund ÅK, Ekman D, Bornberg-Bauer E, Elofsson A: Arrangements in the modular evolution of proteins. Trends Biochem Sci 2008, 33:444–451.

34. Zhong Y, Cheng Z-MM: A unique RPW8-encoding class of genes that originated in early land plants and evolved through domain fission, fusion, and duplication. Sci Rep 2016, 6:32923.

35. Bomblies K, Lempe J, Epple P, Warthmann N, Lanz C, Dangl JL, Weigel D: Autoimmune response as a mechanism for a Dobzhansky-Muller-type incompatibility syndrome in plants. PLoS Biol 2007, 5:e236.

36. Chae E, Bomblies K, Kim ST, Karelina D, Zaidem M, Ossowski S, Martin-Pizarro C, Laitinen RA, Rowan BA, Tenenboim H, et al: Species-wide genetic incompatibility analysis identifies immune genes as hot spots of deleterious epistasis. Cell 2014, 159:1341–1351.

37. Giolai M, Paajanen P, Verweij W, Witek K, Jones JDG, Clark MD: Comparative analysis of targeted long read sequencing approaches for characterization of a plant’s immune receptor repertoire. BMC Genomics 2017, 18:564.

38. Dangl JL, Horvath DM, Staskawicz BJ: Pivoting the plant immune system from dissection to deployment. Science 2013, 341:746–751.

39. Ellis JG, Lagudah ES, Spielmeyer W, Dodds PN: The past, present and future of breeding rust resistant wheat. Front Plant Sci 2014, 5:641.

40. Löytynoja A, Goldman N: A model of evolution and structure for multiple sequence alignment. Philos Trans R Soc Lond, Ser B: Biol Sci 2008, 363:3913–3919.

41. Mistry J, Finn RD, Eddy SR, Bateman A, Punta M: Challenges in homology search: HMMER3 and convergent evolution of coiled-coil regions. Nucleic Acids Res 2013, 41.

42. Stamatakis A: RAxML version 8: A tool for phylogenetic analysis and post-analysis of large phylogenies. Bioinformatics 2014, 30:1312–1313.

43. Letunic I, Bork P: Interactive tree of life (iTOL) v3: an online tool for the display and annotation of phylogenetic and other trees. Nucleic Acids Res 2016, 44:W242–245.

44. Bailey TL, Boden M, Buske FA, Frith M, Grant CE, Clementi L, Ren J, Li WW, Noble WS: MEME SUITE: tools for motif discovery and searching. Nucleic Acids Res 2009, 37:W202–208.

45. Edgar RC: MUSCLE: multiple sequence alignment with high accuracy and high throughput. Nucleic Acids Res 2004, 32:1792–1797.

46. Quinlan AR, Hall IM: BEDTools: a flexible suite of utilities for comparing genomic features. Bioinformatics 2010, 26:841–842.

47. Krzywinski M, Schein J, Birol I, Connors J, Gascoyne R, Horsman D, Jones SJ, Marra MA: Circos: an information aesthetic for comparative genomics. Genome Res 2009, 19:1639-1645.

